# *In-situ* non-lethal rapid test to accurately detect the presence of the nematode parasite, *Anguillicoloides crassus*, in European eel, *Anguilla anguilla*

**DOI:** 10.1101/2020.06.16.155002

**Authors:** M. De Noia, R. Poole, J. Kaufmann, C. Waters, C. Adams, P. McGinnity, M. Llewellyn

## Abstract

*Anguillicoloides crassus* is an invasive nematode parasite of the European eel, *Anguilla anguilla*, and one of the primary drivers of eel population collapse. The presence of the parasite has been shown to impact many features of eel physiology and life history. Early detection of the parasite is vital to limit the spread of *A. crassus*. However, until recently, accurate diagnosis of infection could only be achieved via terminal dissection. To support *A. anguilla* fisheries management in the context of *A. crassus* we developed a rapid non-lethal and non-invasive environmental DNA method to detect the presence of the parasite in the swim bladder. Screening of 131 wild eels was undertaken between 2017 and 2019 Ireland and UK to validate the procedure. DNA extractions and PCR were conducted using both a Qiagen Stool kit at Glasgow University and in situ using Whatman qualitative filter paper No. 1 and a miniPCR DNA Discovery System™. Primers were specifically designed from the cytochrome oxidase mtDNA gene region. In situ extraction and amplification takes approx. 3h for up to 16 individuals with higher specificity and sensitivity compare to the laboratory Qiagen kit extraction. The local diagnostic procedure demonstrated Positive Predictive Values at 96% and Negative Predictive Values at 87%. Our method will be a powerful tool in the hands of fisheries managers to help protect this iconic but critically endangered species. It will allow a non-invasive monitoring of the *A. crassus* dispersion across the European waters.

## 1. Introduction

*Anguillicoloides crassus* (Kuwahara, Niimi & Itagaki 1974) is a nematode is a natural parasite of the *Anguilla japonica* but his final host are Anguilla spp., as the European eel, *Anguilla anguilla* (Lefebvre et al. 2012). The parasite originates from East Asia, having been introduced into Europe in the early 1980s as a result of the trade in live Japanese eels (*Anguilla japonica)* (Weclawski et al. 2013, Laetsch et al. 2012). *A. crassus* is now well established in the Western Hemisphere and can be found in almost all European rivers and lakes, where it can tolerate salinities up to 12 ppt (Aguilar et al. 2005, Becerra-Jurado et al. 2014). *A. crassus* has been identified as one of the main contributing factors to the collapse of eel populations in Europe (Henderson et al. 2012).

*A. crassus* reproduces sexually in the swim bladder of the eels. The eggs hatch in the female worm inside the swim bladder and L2 larvae migrate to the intestinal tract via blood vessels to be excreted with the faeces (Didžiulis, 2013). As part of its life cycle *A. crassus* is then trophically transmitted to various intermediate hosts including several zooplankton species (especially copepods of the orders Cyclopoidea and Calanoidea) as well as small fish such as the three-spined sticklebacks, *Gasterosteus aculeatus* (Kuwahara, 1999). In the intermediate host, the parasite develops into the infectious phase L4 larvae, which, once ingested, parasitize the eel as the final host (IUCN 2013). Parasite migrate from the gut system perforating the connective tissue and muscles reaching the swim bladder (Heitlinger et al, 2009). The number of parasites found in the swim bladder can vary from less than ten to greater than 70 individuals per eel. The presence of the parasite has been shown to detrimentally affect many features of eel physiology and life history (Newbold et al., 2015). Adult nematodes feed on blood supplied to the swim bladder wall and can result in increased eel mortality as a consequence of damage caused to the organ (Schneebauer et al. 2017). The swim bladder wall becomes thinner and less elastic due to the perforation caused by the parasite feeding habit with an impacting buoyancy control (Weclawski et al., 2013, Newbold et al., 2015, Barry et al., 2014) *A. crassus* infection is also thought to alter the physiological mechanisms involved in silvering – the process by which freshwater sub-adults adapt to life in the ocean. In this respect, infected eels have also been found to silver faster as a result of an over production of cortisol, which seems to have a stimulatory effect on GTH2 synthesis (Di Biase et al., 2017). Moreover, cortisol is the key hormone produced during fasting, typical of the silvering phase stage (Fazio et al. 2012). Due to their blood feeding behaviour, the parasites increase erythropoiesis in infected eels, which is a trait that normally increases during silvering (Churcher et al., 2015). The presence of the parasite can also affect the eel’s migratory speed and behaviour during the migration to the Sargasso sea (Pelster 2015, Newbold et al. 2015). It has been proposed by Righton et al. (2016) that the eel’s migration speed in the ocean is reduced and the energy demand increases, due to the reduction of the swim bladder elasticity. The presence of the parasite appears not to affect the speed and migratory behaviour during the first phase of the migration in shallow water (Simon et al, 2018). However, where deep diving is required in the ocean, damage to the integrity of the swim bladder is believed to seriously impact on an infected eel’s chances of survival (Righton et al. 2016).

Accurate detection of the parasite can only currently be achieved via post-mortem dissection and thus requires the eel to sacrificed. However, several non-lethal techniques are under development (Frisch, Davie, Schwarz, & Turnbull, 2016). Anal redness can be used as indicator for presence or absence of the parasite, but this approach lacks both specificity and objectivity (Crean et al. 2003). A radio diagnostic method has been developed to detect inflammation caused by the nematode’s feeding habits (Beregi, Molnár, Békési, & Székely, 1998). The method uses X-ray to scan the pneumatic duct and can detect swim bladder damage and parasite presence. The quality of the images has a large margin of error so the accuracy of detection can be low and swim bladder alterations can be caused by other factors (Beregi et al., 1998). Frisch et al. 2016 made improvements to the method developed by Beregi et al. (1998). Using compound radiography they were able to detect small alterations to the thickness of the swim bladder wall and to inflations of the lumen. However to perform a full body scan using this method, the animal has also to be euthanized.

To support *A. anguilla* fisheries management in the context of *A. crassus*, rapid non-lethal and non-invasive methods for pathogen detection are urgently required. Screening of translocated eel populations could, for example, limit the spread of the pathogen. Furthermore, non-lethal screening of silver eels alongside satellite tagging studies could reveal the impact of infection on the migratory and breeding success. Non-invasive molecular based methods have been proposed to detect parasites in various fish species related to the food health safety chain and conservation management (Levsen et al. 2018, Cavallero et al. 2017). In other studies quantitative qPCR has been developed to detect *Anisakis* sp. in experimentally prepared fish-derived food products (Godínez-González et al. 2017, Mossali et al. 2010). A non-lethal qPCR based eDNA approach has also been optimized to detect the cestode *Schistocephalus solidus* in samples taken by needle from the intra-peritoneal cavity of a fish (Chloé Suzanne Berger & Aubin-Horth, 2018).

In the current study, we developed a non-invasive, non-lethal and portable, *in situ* PCR-based approach to detect *A. crassus* in the European eel. We tested the approach’s specificity and sensitivity at field sites in the UK and Ireland, as well as exploring the link between host condition and parasite infection status.

## 2. Materials and Methods

### 2.1 Sample collection

The study was conducted at two different locations in UK and Ireland. In the Burrishoole catchment, Ireland 53°55‘27.6“N 9°34‘27.0“W, yellow were collected from Loughs Feeagh and Furnace using unbaited fyke nets deployed overnight in chains of 10 nets sets at different lake depths in spring 2017, 2018 and 2019. Silver eels were collected in autumn 2019 using river traps (Poole, Rogan, & Mullen, 2007). The study was carried out under a Health Products Regulatory Authority (HPRA) license number AE19130-P096. Laugh Feeagh in a fresh water lake and Lough Furnace is a brackish water lake. In Lough Neagh, UK-Northern Ireland 54°36‘05.5“N 6°24‘55.5“W, yellow eels were collected with baited long lines fished in the lake during the morning in summer 2018. The eels were euthanized with an overdose of MS-222 followed by a cervical separation of the spinal cord. Fat content was measured using a Distell Fatmeter Model - FM 692. For each eels weight and length was recorded (Supplementary 1). A colonic irrigation with 2ml 0.09 % sterile saline solution was performed to collect faecal material using a 5 ml syringe and a Terumo Surflo Winged Infusion Set 23g Continuous Use with the needle removed (Figure 1). A drop of the extracted wash material was placed on 1 cm^2^ of Whatman qualitative filter paper No. 1 and air dried for 5 minutes at room temperature, then stored at −80°C. The remaining wash was stored in 100% ethanol (1 wash:9 ETOH) at −20°C. Subsequently, eels were dissected, swim bladder inspected and the number of *A. crassus* present counted. *A. crassus* were collected and stored in 100% ethanol before being stored at −20°C.

**Fig. 1.**
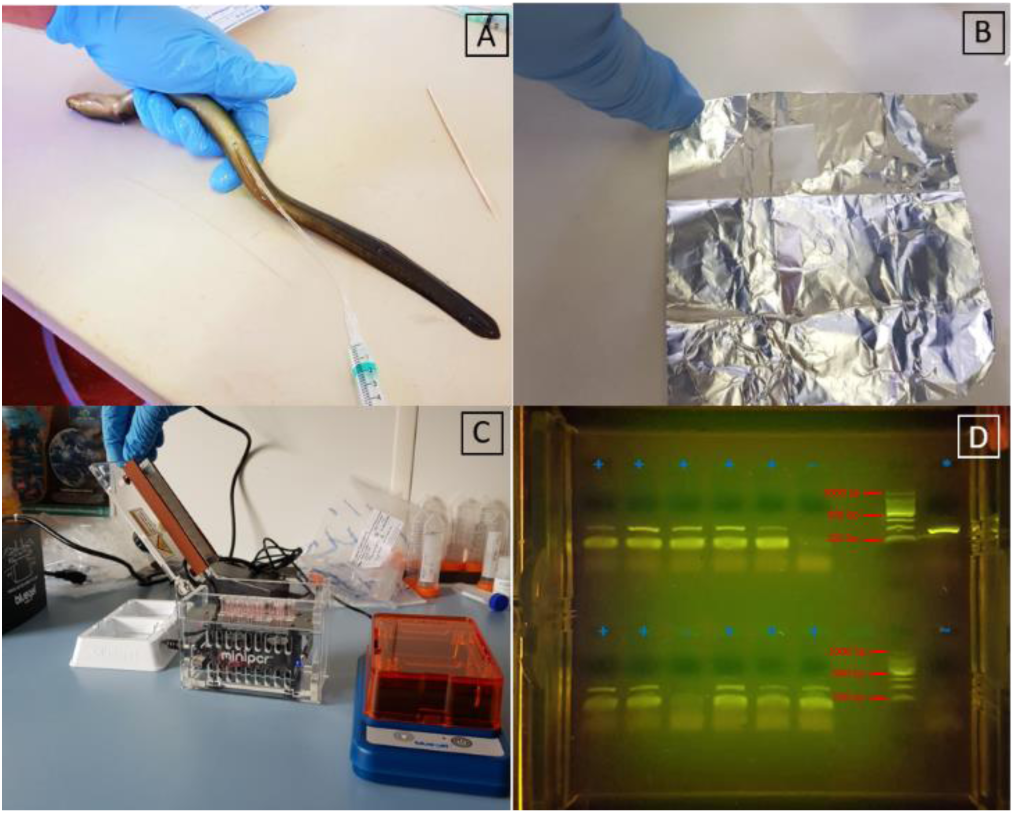
A) Colonic irrigation with sterile saline solution (9 ‰) on an anesthetized yellow eel. B) Collection of a drop of faecal material on a piece of Whatman qualitative filter paper No. 1. C) *In situ* DNA extraction and diagnostic PCR with MiniPCR thermocycler. D) *In situ* visualization on electrophoresis agarose gel 2 % on amplified target CO1 gene. “+” Positive amplification from faecal extracted DNA, “-” Negative amplification from faecal extracted DNA, “*” Positive control, “∼” Negative control.

### 2.2 DNA collection and extraction methods development

#### 2.2.1 Laboratory genomic DNA extraction

DNA from 200 µl of stored faecal material was extracted with the QIAamp Stool Kit (Qiagen) with a slight modification to the Qiagen Stool Kit suggested protocol. ATL tissue lysis buffer volume was increased to 350 µl, proteinase K up to 20 µl, AL lysis buffer up to 300ul and Ethanol 100% up to 400ul. All quantities refer to individual samples.

#### 2.2.2 In situ genomic DNA extraction

A small sample of the filter paper of 1mm diameter was removed by a punch from the Whatman qualitative filter paper No. 1 were faecal material had been previously deposited and DNA was extracted using adjusted extraction protocol of DNA from Whatman™ FTA™ cards (Santos, Sasal, Verneau, & Lenfant, 2006) (Supplementary 1) (Figure 1).

### 2.3 Primer design, PCR conditions and species specificity

A pair of specific customised primers were designed using the most conserved region within the ten cytochrome c oxidase subunit 1 (COI) gene sequences available for *A. crassus* (NCBI Accessions: MF458547). The sequences of *A. crassus* COI were aligned to identify a conserved region within *A. crassus* suitable for primer design (Table 1). The obtained target sequence was blasted against COI sequences for three of the most common fish nematode parasites *Camallanus* sp. (NCBI Accession: EU598889), *Contracaecum* sp. (NCBI Accession: FJ866816) and *Capillaria* sp (NCBI Accession: AJ288168). to confirm the unique target region in *A. crassus* (Pouder, Curtis, & Yanong, 2009).

**Table 1.**
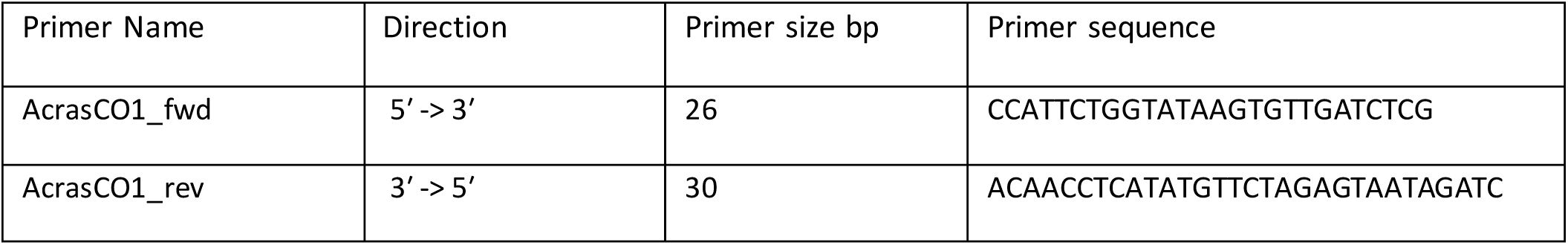
Primer name, direction of amplification, primer size expressed in base pairs and specific designed sequence.

The total volume of the PCR reaction was 20 µl. The cycle used for the PCR started with 5 minutes at 95 °C, followed by 35 cycles of 95 °C for 30 seconds, 60 °C for 30 seconds and 72 °C for 30 seconds and a last step of 10 minutes at 72 °C. The PCR products were visualized on a 2% agarose gel using SYBR safe staining (Invitrogen). Species specificity of the primer set was tested using a series of nematode parasites collected in the same study system, various non-nematode parasites and other animal taxa, including the European eel, to assay cross-reactivity (*Anisakis* sp., *Lepeophtheirus salmonis, Anguilla anguilla, Paramoeba perurans, Scombrus scombrus, Diphyllobothrium* sp., *Schistocephalus* sp., *Dentitruncus truttae*).

### 2.4 Specificity, Sensitivity NPV and PPV

Several parameters were calculated to assay the validity of the test. Here sensitivity is the ability of a test to correctly classify an individual as infected. The ability of a test to correctly classify an individual as non-infected is called the test′s specificity. The Positive Predictive Value (PPV) is the percentage of eels with a positive test which are actually infected and the Negative Predictive Value (NPV) is the percentage of eels with a negative test which do not have the parasite. Positive and negative predictive values are directly related to the prevalence of the disease in the population (Stojanović et al., 2014) (Table2).

**Table 2.**
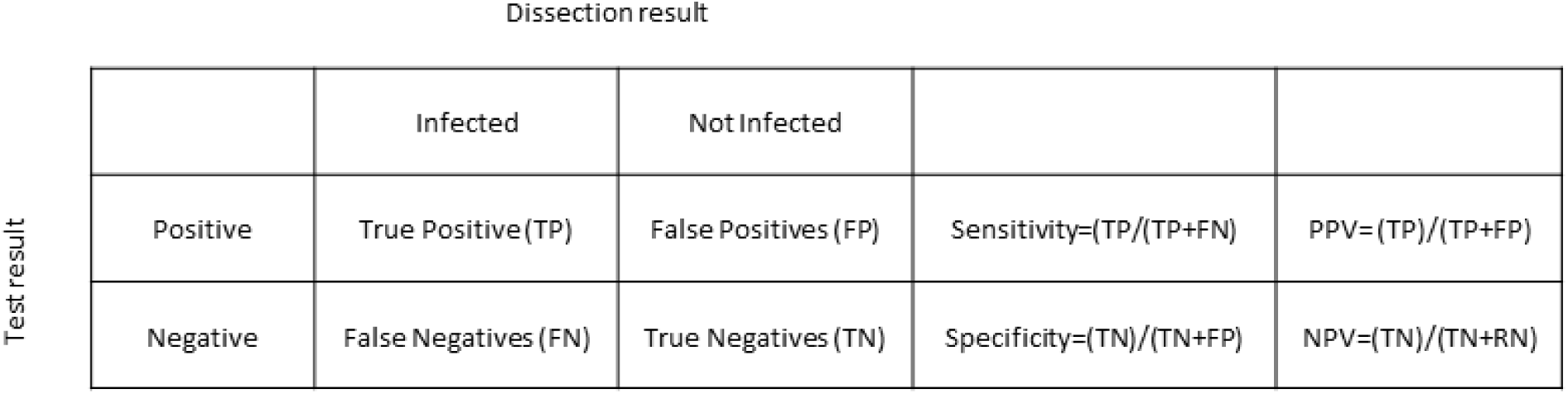
The criteria for Specificity, Sensitivity NPV (Negative Predictive Value) and PPV (Positive Predictive Value). Animals that are infected and test as positive are considered True Positive (TP). Animals are infected and test negative are described as False Negative (FN). Animals that have no infection, but test positive are False Positive (FP), and those not infected but test negative are True Negative (TN).

### 2.5 Biological validation

#### 2.5.1 Eggs count in faecal wash

Nematode larvated and unlarvated eggs and L2 larvae were counted with a modified McMaster Salt Flotation Technique. 200 µl of faecal material was diluted in 1,5 ml of distilled water. After mechanical homogenisation, the suspension was poured through a 250 micron aperture sieve and the filtrate collected. After thorough mixing, the solution was transferred to a centrifuge tube and spun for five minutes at 2500 rpm. The supernatant was discarded and the remaining faecal pellet covered and homogenised with 300 µl of saturated sodium chloride solution, mixed by inverting slowly six times. Then, using a Pasteur pipette, the mixture was transferred to a McMaster slide. Each chamber holds 0.15 ml beneath the gridded area. The preparation is then examined using the x 25 objective of a stereo microscope, the number of eggs present in the grids of both chambers were counted to give an estimate of the numbers of eggs/gram of faecal material.

#### 2.5.2 Nematode count in swim bladder

All dissected eels were checked for *A. crassus*, and where present, they were counted. The swim bladder was extracted whole from the animal and stored in the fridge till the procedure was completed for all the specimens. The swim bladder was then opened and nematodes were counted and classified to adults and larval stages.

### 2.6 Statistical analysis

We used a generalized linear mixed model (GLMM) using environmental factors and eel condition phenotypes as variables to predict the eel parasite load and prevalence. Eel identity was considered as a random factor. Length, weight, year of sampling, condition factor, fat and salinity were considered as explanatory variables. Correlation between variables was assessed using likelihood-ratio tests (ANOVA). The best fitting model was selected using dredge MuMIn package in R using Akaike’s Information Criterion (AICc), Bayesian Information Criterion (BIC) and log-likelihood for selection criteria with package TmB in R.

## Results

### 3.1 DNA extraction and primers Specificity

A total of 131 eels were collected over the three sampling years. DNA was extracted from 104 fish using the Qiagen Stool Kit. The Whatman extraction protocol was used on 55 fish, with an overlap of 28 extractions were both methods were deployed. DNA was successfully extracted from all samples using both Qiagen and Whatman methods. Primers specificity was confirmed when amplification in different species of fish parasite nematodes and various other species failed, not presenting a positive band in the PCR amplification product. Confirmation of the specific amplification was obtained from sequences obtained using single end Sanger-purified PCR products.

### 3.2 Eggs count in faecal material

Faecal material from 131 samples was tested to detect the presence of eggs and/or L2 larvae using the McMaster floatation protocol. Samples were analysed in triplicates. No eggs or larvae were detected.

### 3.3 Rapid test *in-situ*

An *in situ* non-invasive test was performed on 55 eels collected in 2019. Individuals were rectally catheterised to enable colonic irrigation without causing internal lesions. The amount of saline solution used varied according to the size of the tested animals. Smaller eels required a smaller volume of saline solution to not overflood the intestine. The drop of collected material was air dried at RT on Whatman paper. After drying the paper was used for extraction or stored −80 C. *In-situ* DNA extraction took 20 minutes for 16 samples, PCR reaction was undertaken over a period of hours and electrophoresis with gel visualization took a further 17 minutes. Thus, the test can be performed for 16 samples in approximately 3 hours. The cost per eel for each test is £2 and the cost of the associated equipment setup approximately £900 (Figure 1).

### 3.4 Comparison of different DNA extraction methods

The DNA Whatman paper extraction method provides a rapid and more reliable assessment of infection (sensitivity and specificity) compared to the method based on the Qiagen Stool and Blood kit. Using the Whatman paper, the time for DNA extraction from 1 sample is reduced from 1 hour to 20 minutes and cost from £3.70 to £2 per sample, with possibility of running 16 samples in parallel. Even though the number of eels tested with the Whatman qualitative filter paper No. 1 DNA extraction method was lower than those using the Qiagen approach, (55 compared to 104), there was an overall reduction of false positive and false negative PCR amplification results (Table3) (Figure 2).

**Table 3.**
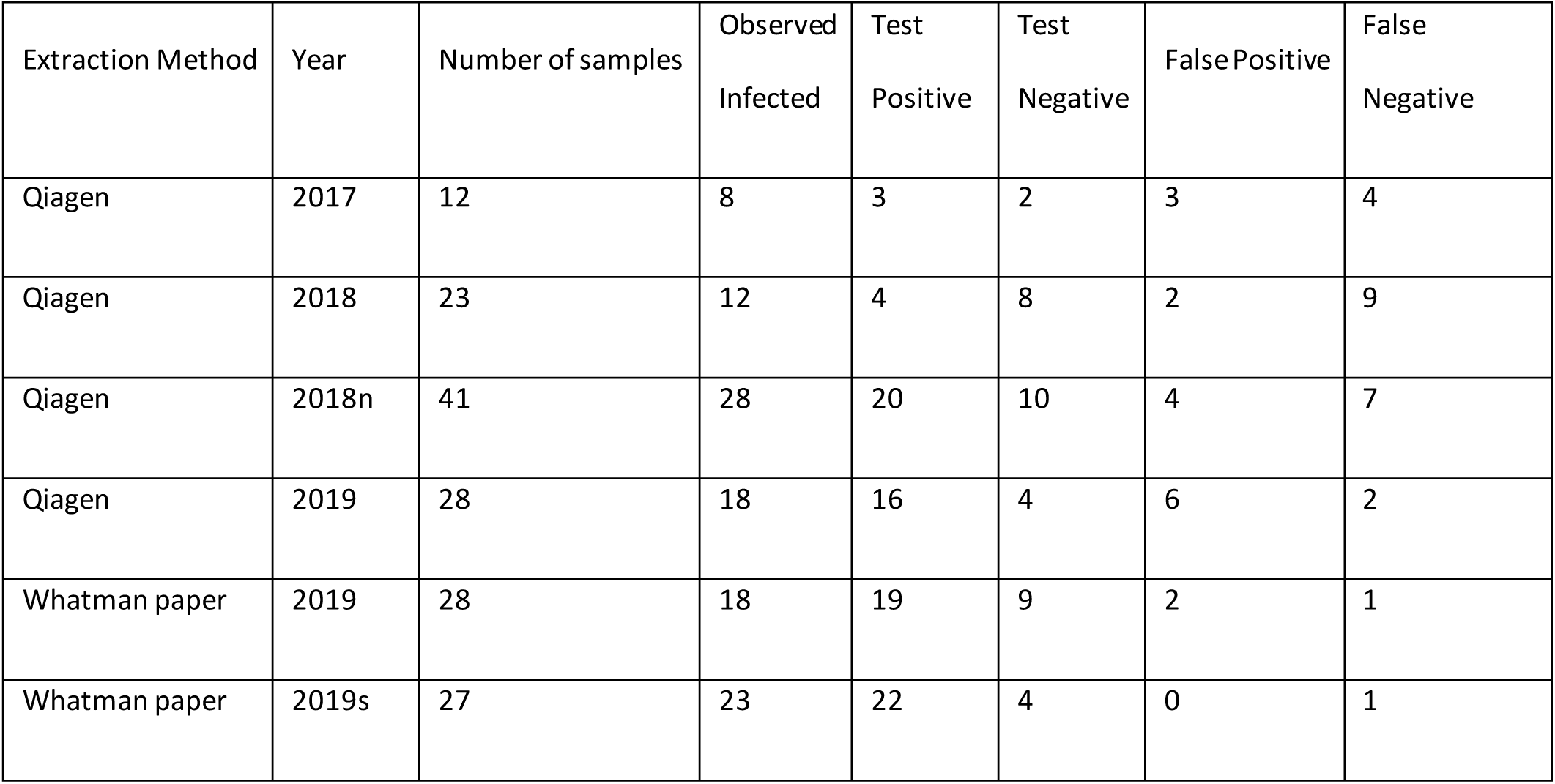
Relative test detection for the two different extractions methods across the 3 sampling seasons. For 2019 the same samples were tested with both methods. 2018n considers samples collected in Lough Neagh and 2019s silver eels collected in the Burrishoole catchment.

**Fig. 2.**
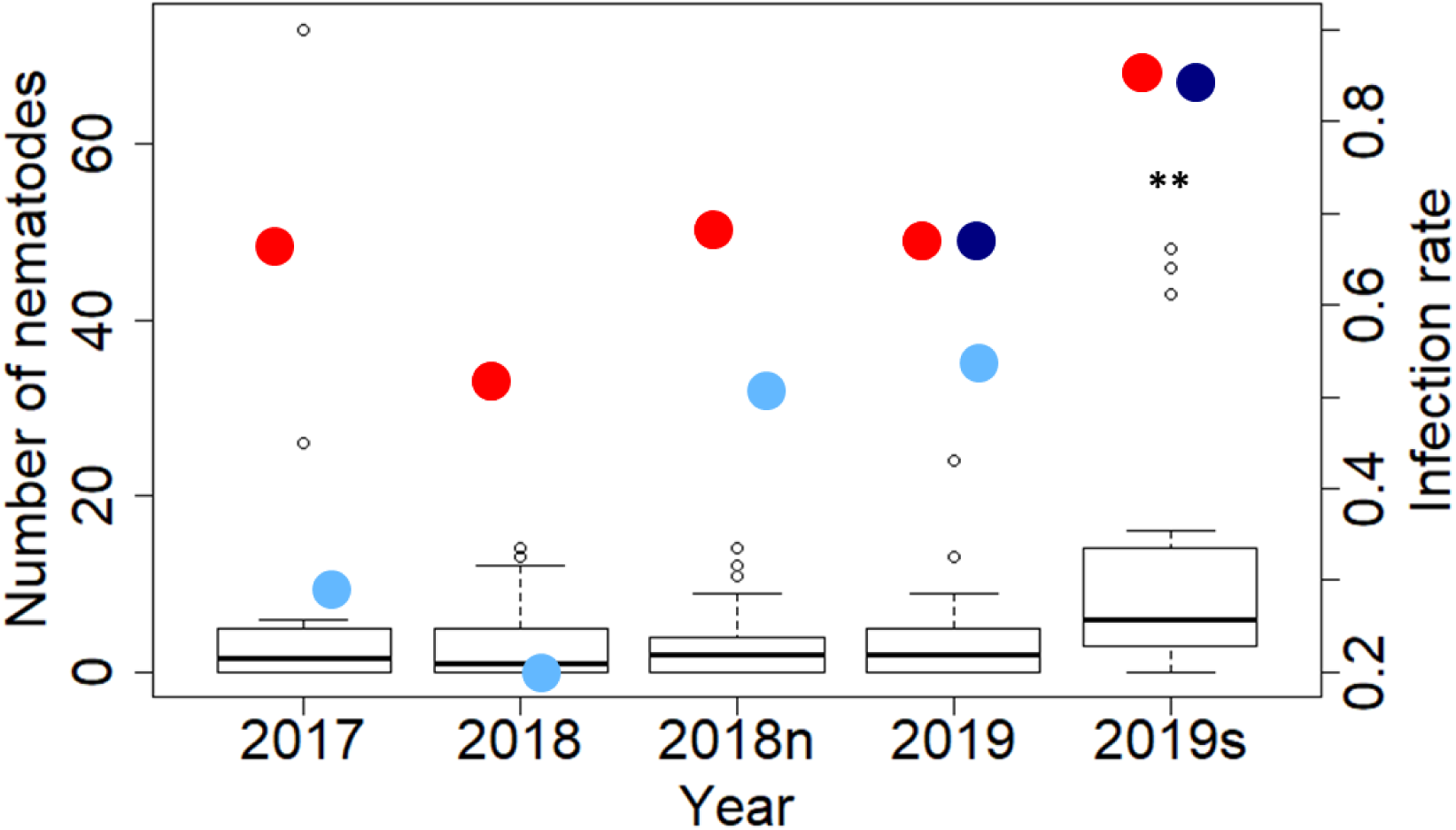
Number of *Anguilicola crassus* counted in dissected animals and infection rate in different years of sampling. Infection rate represents the number of animal infected compared to the total number of animals. Red dots show the actual infection rate based on counts acquired from dissected animals. Light Blue dots indicate the infection rate derived from the extraction using Qiagen Blood and Stool kit. Dark Blue dots indicates the infection rate observed with the rapid test. A Mann-Whitney test shows that silver eels in 2019 were significantly more infected than in the previous year (P value < 0.05). “2018n” refers to samples collected in Lough Neigh and “2019s” to silver eels collected in the Burrishoole system.

### 3.5 Specificity/Sensitivity/NPV/PPV

The Whatman rapid test returns a higher proportion of PPV and NPV detections. The resulting improvement of using the Whatman test in specificity is 46%, in sensitivity 45%, in PPV is 30% and in NPV is 41% (Figure 3).

**Fig. 3.**
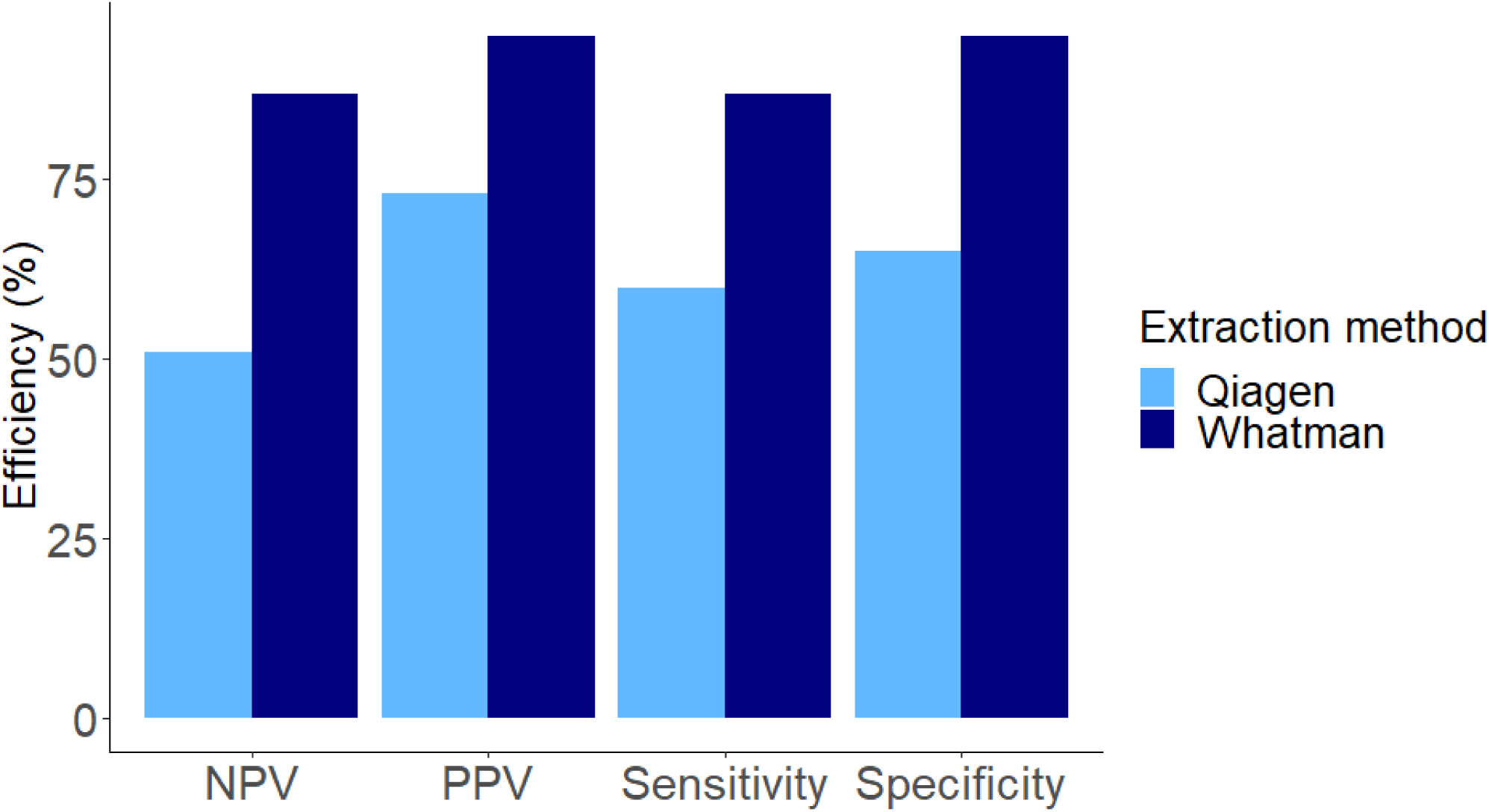
Relative parasite detection efficiency for the two DNA extraction methods. The Whatman DNA extraction method performs better in all the categories with an average improvement of 41% over the Qiagen method.

### 3.6 Statistical analysis of *A. crassus* infection covariates

Statistical analysis to explore the drivers of infection prevalence was undertaken for samples collected in the Burrishoole system. Among all the variables taken into consideration for the model only Lake, Year of collection, Fat and Lake were identified on the basis they showed no signs of collinearity. The GLMM that best fitted the data was a zero inflation model with fat and length as the most reliable predictors the parasitic load (AIC=488 and LolLi= −236 with p-value <0.001). Fat best explained infection prevalence with a quadratic function with the highest infection rate for those of around average fat content Fat was corrected with a quadratic factor after tested with ANOVA, with length kept as a linear factor (Figure 4.). Parasitic load was found to increase with the length of the animal. The most infected eels were those with an average fat content.

**Fig. 4.**
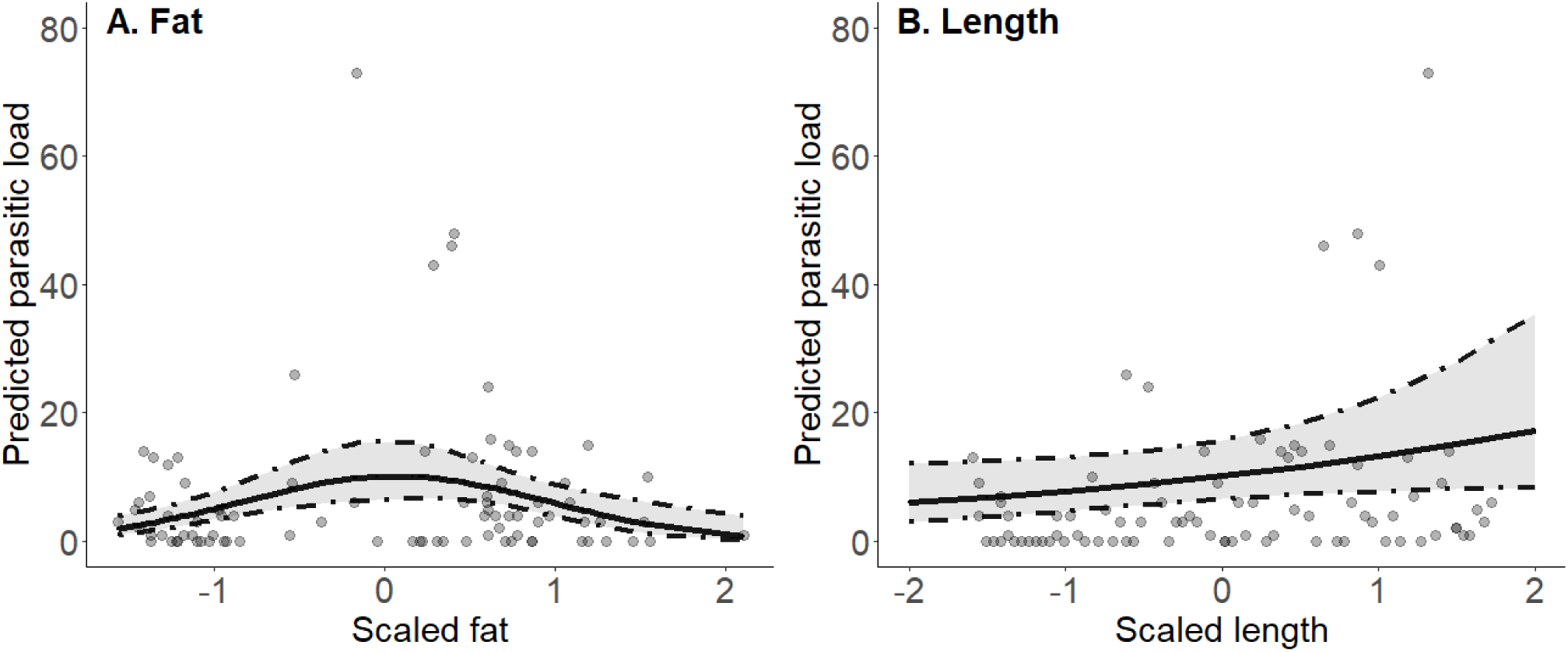
Observed data super-imposed on the model-predicted *A. crassus* parasitic load. The lines represent model predictions and the points represent experimental data. On x axis scaled fat and length for graphic purpose. The parasite load is predicted to increase with longer animals and with animals recorded with a mean fat content.

## Discussion

Our study represents the first attempt to develop a sensitive, non-lethal and i*n situ* method to establish *A. crassus* infection in *A. anguilla* via the detection of parasite-shed eDNA in faecal material. High values for NPV (87%) and PPV (95%) suggest that the tool may have a role in both veterinary control and fisheries management contexts. We found inter-annual differences in the number of infected eels in the three years we sampled in the Burrishoole catchment with the number of infected animals across the sampling seasons. Furthermore, we noted a clear correlation between parasite burden and physical characteristics of infected individuals. Highly infected when dissected were showing a damaged swim bladder, with reduction of the lumen and thickening of the wall, as suggested by Frisch et al. (2016).

Mitochondrial genes are the major target genes in PCR-based detection systems because they are highly conserved and present in multiple copies (Paoletti et al., 2018). High target copy number may explain the high sensitivity of the test we deployed. The mitochondrial gene COI, has been widely used to detect the presence of nematode parasites in commercially important fish species (Godínez-González et al., 2017; Herrero, Vieites, & Espiñeira, 2011; Paoletti et al., 2018; Santos et al., 2006). Many studies rely on qPCR or Real Time PCR for fish pathogens detection, but such approaches are challenging and expensive to deploy in the field (Chlo Suzanne Berger & Aubin-Horth, 2018; Hutchins et al. 2018; Trujillo-González et al. 2019).

Intriguingly, a major improvement in detection sensitivity and specificity was achieved here by adopting a more ‘crude’ nucleic acid extraction approach (Figure 3). Nucleic acid extraction is increasingly recognised as a major rate limiting step in molecular diagnostics, however, paper-based options offer several advantages in terms of speed and cost (Zou et al., 2017). The increased sensitivity of our paper-based approach over column-based kits likely stems from the higher concentrations of DNA that are recovered.

Our inability to microscopically detect the presence of *A. crassus* larvae or eggs in faecal material, suggests DNA or cellular fragments from worms are the likely source of the genetic material we detected. DNA-based molecular diagnostics for helminth parasite detection are gaining in popularity (O’Connell & Nutman, 2016). However, as has been observed for the detection for many pathogens of medical and veterinary interest, qPCR is preferred over standard PCR to deliver the sensitivity required (Thomas et al., 2019). Point-of-care qPCR for viral pathogens can now deliver a result in less than 20 minutes (Melchers et al, 2017). Similarly, several mobile qPCR instruments have been brought to the market and have been successfully deployed to deliver veterinary diagnoses in remote locations on actionable timescales (Hole & Nfon, 2019). However, the cost of such approaches is likely to be prohibitive in respect to their application to the detection of pathogens in non-human subjects. Our approach, validated by dissection and exhaustive worm counts, shows that standard PCR, using low cost reagents and equipment may be just as portable, and informative epidemiologically, as ‘higher-end’ devices. Our test fits in a small suitcase and can be performed in the field powered with a motorcycle battery. Stocking as part of eel population enhancement is the major contributor to *A. crassus* dispersal (Weclawski et al. 2013, Laetsch et al. 2012). Early detection with a non-lethal method could be a powerful tool to avoid the spread of the parasite and monitoring purpose.

*A. crassus* has multiple effects on the eels, from a physiological, immunological and pathological point of view (Laetsch et al., 2012). Post infection the swim bladder becomes increasingly thickened and opaque as a result of fibrosis (Kirk, 2003). The blood-feeding activities of *A. crassus* cause degenerative and inflammatory changes compromising the buoyancy of the animal during their fresh water stage, but more importantly during their ocean migration to the spawning grounds (Righton et al., 2016). Our GLMM shows how the length of the eels is related to the level of infection with higher levels of parasites in larger fish. Longer animals are mostly older animals and they have more chances to encounter the parasite (Weclawski et al., 2013). The interpretation of the correlation between fat and infection rate is unclear. Eels entering the catchment are low in fat and present low infection rate because they have not encountered the parasite in their saltwater stage; *A. crassus* has not been found in the marine environment. When they fatten during the yellow stage of the life cycle the possibility of being infected increases as their fat level increases. Peculiar is the reduction of the infection in eels with high levels of fat, this needs to be investigated further.

The increase in parasite load we noted from 2017 to 2019 follows a trend that is also found all over Europe, where the parasite is established and is fast colonizing all the fresh water basins (Aguilar et al., 2005; Schabuss, Kennedy et al., 2005; Selim & El-ashram, 2012; Wielgoss et al. 2008). *A. crassus* has a recent history in the Burrishoole catchment. First detection occurred in 2010 in a yellow eel and in 2016 for the first time in a silver eel. In contrast to the rising burdens across much of Europe, in some lakes where the parasite had been detected since first discovery there is stabilization and even a slight decline in nematode abundance and intensities (Wielgoss et al., 2008). There is a possibility of an increased resistance towards the parasites in the long term (Schabuss et al., 2005). Although some evidence of increasing tolerance of *A. anguilla* to parasite infection, the overall impact of the parasite on the eel’s mortality has been severe and is likely a contributor to the European population’s steep decline impeding a recovery (Kirk, 2003; Schabuss et al., 2005). Treatment of infected eels with anti-helminthic has not been trialled and a single study into a vaccination test to reduce the development of adult from irradiated L3 larvae showed that the antibody response is not a key element in resistance of eels against *A. crassus* (Knopf & Lucius, 2008). Infection control, which requires extensive diagnostic testing, therefore represents the only feasible route to reducing population-wide parasite burden.

In this study, we developed a powerful tool that can be used for specific parasite screening for the European eel. Cost per sample is low and the time to run a test comprising 16 samples is under 3 hours. Future work should focus on the improvement of the specificity. Our test offers managers the opportunity to engage in efficient infection control by assessing the disease status of eels before allowing transfers between river systems. The rapid test represents an important contribution to the conservation and management of this critically endangered species.

## Acknowledgment

This research was supported in part by a research grant from the BBSRC (grant number BB/P001203/1), by Science Foundation Ireland, the Marine Institute, Ireland, and the Department for the Economy, Northern Ireland, under the Investigators Program grant number SFI/15/IA/3028. We would like to express our sincere gratitude to Dr Derek Evans to provide the samples from Lough Neagh, to Sara Gandy for the support in statistical analysis and to all the member of the Marine Institute in Newport, Ireland, for the immense help during sampling.

