## Supplementary 1 for "*In-situ* non-lethal rapid test to accurately detect the presence of the nematode parasite, *Anguillicoloides crassus*, in European eel, *Anguilla anguilla*"

Supplementary 2. Data collection

| **Eel** | **Year of collection** | **Location** | **Nematode counted** | **Infection rate** | **Extraction method** | **Life stage** | **Length (cm)** | **Weight (g)** | **Fat** |
| --- | --- | --- | --- | --- | --- | --- | --- | --- | --- |
| 1 | 2017 | Burrishoole | 0 | 0.67 | Qiagen | Yellow | 31.3 | 55 | N/A |
| 2 | 2017 | Burrishoole | 26 | 0.67 | Qiagen | Yellow | 38.3 | 90 | 12.5 |
| 3 | 2017 | Burrishoole | 6 | 0.67 | Qiagen | Yellow | 40 | 105 | 15.4 |
| 4 | 2017 | Burrishoole | 1 | 0.67 | Qiagen | Yellow | 42.3 | 140 | 7.6 |
| 5 | 2017 | Burrishoole | 1 | 0.67 | Qiagen | Yellow | 36.6 | 90 | 12.3 |
| 6 | 2017 | Burrishoole | 73 | 0.67 | Qiagen | Yellow | 53.9 | 275 | 15.5 |
| 7 | 2017 | Burrishoole | 4 | 0.67 | Qiagen | Yellow | 47.6 | 185 | 21.7 |
| 8 | 2017 | Burrishoole | 4 | 0.67 | Qiagen | Yellow | 36.2 | 70 | 9.6 |
| 9 | 2017 | Burrishoole | 0 | 0.67 | Qiagen | Yellow | 36.7 | 75 | 9.1 |
| 10 | 2017 | Burrishoole | 2 | 0.67 | Qiagen | Yellow | 58.1 | 320 | 5 |
| 11 | 2017 | Burrishoole | 0 | 0.67 | Qiagen | Yellow | 58.1 | 320 | 5 |
| 12 | 2017 | Burrishoole | 0 | 0.67 | Qiagen | Yellow | 36.6 | 90 | 12.3 |
| 13 | 2018 | Burrishoole | 0 | 0.52 | Qiagen | Yellow | 49 | 200 | 6.6 |
| 14 | 2018 | Burrishoole | 0 | 0.52 | Qiagen | Yellow | 43.2 | 135 | 6.9 |
| 15 | 2018 | Burrishoole | 5 | 0.52 | Qiagen | Yellow | 37.8 | 90 | 8.9 |
| 16 | 2018 | Burrishoole | 0 | 0.52 | Qiagen | Yellow | 33.7 | 70 | 18.2 |
| 17 | 2018 | Burrishoole | 0 | 0.52 | Qiagen | Yellow | 34.3 | 65 | 20.8 |
| 18 | 2018 | Burrishoole | 6 | 0.52 | Qiagen | Yellow | 33.7 | 65 | 25.8 |
| 19 | 2018 | Burrishoole | 14 | 0.52 | Qiagen | Yellow | 57.3 | 365 | 5.2 |
| 20 | 2018 | Burrishoole | 5 | 0.52 | Qiagen | Yellow | 59.9 | 365 | 4.8 |
| 21 | 2018 | Burrishoole | 12 | 0.52 | Qiagen | Yellow | 50.7 | 205 | 6.4 |
| 22 | 2018 | Burrishoole | 1 | 0.52 | Qiagen | Yellow | 59 | 285 | 5.6 |
| 23 | 2018 | Burrishoole | 0 | 0.52 | Qiagen | Yellow | 47.8 | 150 | 7.5 |
| 24 | 2018 | Burrishoole | 3 | 0.52 | Qiagen | Yellow | 64.2 | 475 | 4 |
| 25 | 2018 | Burrishoole | 0 | 0.52 | Qiagen | Yellow | 36.1 | 90 | 19.7 |
| 26 | 2018 | Burrishoole | 0 | 0.52 | Qiagen | Yellow | 35.4 | 90 | 16.5 |
| 27 | 2018 | Burrishoole | 9 | 0.52 | Qiagen | Yellow | 38.2 | 100 | 27.2 |
| 28 | 2018 | Burrishoole | 1 | 0.52 | Qiagen | Yellow | 36.2 | 90 | 22.8 |
| 29 | 2018 | Burrishoole | 3 | 0.52 | Qiagen | Yellow | 37.2 | 90 | 20.9 |
| 30 | 2018 | Burrishoole | 0 | 0.52 | Qiagen | Yellow | 96.1 | 266 | 18.7 |
| 31 | 2018 | Burrishoole | 0 | 0.52 | Qiagen | Yellow | 55.8 | 285 | 7.2 |
| 32 | 2018 | Burrishoole | 1 | 0.52 | Qiagen | Yellow | 46.5 | 230 | 23.3 |
| 33 | 2018 | Burrishoole | 0 | 0.52 | Qiagen | Yellow | 51.1 | 235 | 29.4 |
| 34 | 2018 | Burrishoole | 0 | 0.52 | Qiagen | Yellow | 42.4 | 135 | 18.5 |
| 35 | 2018 | Burrishoole | 13 | 0.52 | Qiagen | Yellow |  |  | 8.2 |
| 36 | 2019 | Burrishoole | 7 | 0.67 | Whatman and Qiagen | Yellow | 52.4 | 245 | 5.5 |
| 37 | 2019 | Burrishoole | 3 | 0.67 | Whatman and Qiagen | Yellow | 41.3 | 105 | 13.8 |
| 38 | 2019 | Burrishoole | 3 | 0.67 | Whatman and Qiagen | Yellow | 45.3 | 190 | 34.2 |
| 39 | 2019 | Burrishoole | 1 | 0.67 | Whatman and Qiagen | Yellow | 41.8 | 115 | 7.8 |
| 40 | 2019 | Burrishoole | 0 | 0.67 | Whatman and Qiagen | Yellow | 51.8 | 160 | 19.4 |
| 41 | 2019 | Burrishoole | 13 | 0.67 | Whatman and Qiagen | Yellow | 31.2 | 50 | 6.9 |
| 42 | 2019 | Burrishoole | 1 | 0.67 | Whatman and Qiagen | Yellow | 34.1 | 75 | 21.9 |
| 43 | 2019 | Burrishoole | 24 | 0.67 | Whatman and Qiagen | Yellow | 39.6 | 90 | 21.9 |
| 44 | 2019 | Burrishoole | 1 | 0.67 | Whatman and Qiagen | Yellow | 58.2 | 265 | 6.1 |
| 45 | 2019 | Burrishoole | 0 | 0.67 | Whatman and Qiagen | Yellow | 51.3 | 260 | 28.9 |
| 46 | 2019 | Burrishoole | 9 | 0.67 | Whatman and Qiagen | Yellow | 39.7 | 100 | 12.4 |
| 47 | 2019 | Burrishoole | 7 | 0.67 | Whatman and Qiagen | Yellow | 33.7 | 75 | 21.8 |
| 48 | 2019 | Burrishoole | 0 | 0.67 | Whatman and Qiagen | Yellow | 32 | 40 | 18.7 |
| 49 | 2019 | Burrishoole | 1 | 0.67 | Whatman and Qiagen | Yellow | 35.9 | 85 | 7.15 |
| 50 | 2019 | Burrishoole | 0 | 0.67 | Whatman and Qiagen | Yellow | 38.4 | 75 | 18.6 |
| 51 | 2019 | Burrishoole | 0 | 0.67 | Whatman and Qiagen | Yellow | 38.3 | 95 | 7.8 |
| 52 | 2019 | Burrishoole | 0 | 0.67 | Whatman and Qiagen | Yellow | 49.4 | 175 | 6.8 |
| 53 | 2019 | Burrishoole | 4 | 0.67 | Whatman and Qiagen | Yellow | 41.7 | 170 | 6.4 |
| 54 | 2019 | Burrishoole | 4 | 0.67 | Whatman and Qiagen | Yellow | 31.5 | 45 | 22.2 |
| 55 | 2019 | Burrishoole | 4 | 0.67 | Whatman and Qiagen | Yellow | 35.9 | 75 | 9 |
| 56 | 2019 | Burrishoole | 0 | 0.67 | Whatman and Qiagen | Yellow | 34.9 | 80 | 26.4 |
| 57 | 2019 | Burrishoole | 4 | 0.67 | Whatman and Qiagen | Yellow | 34.1 | 90 | 24.8 |
| 58 | 2019 | Burrishoole | 0 | 0.67 | Whatman and Qiagen | Yellow | 40.1 | 110 | 8 |
| 59 | 2019 | Burrishoole | 10 | 0.67 | Whatman and Qiagen | Yellow | N/A | N/A | N/A |
| 60 | 2019 | Burrishoole | 2 | 0.67 | Whatman and Qiagen | Yellow | N/A | N/A | N/A |
| 61 | 2019 | Burrishoole | 0 | 0.67 | Whatman and Qiagen | Yellow | N/A | N/A | N/A |
| 62 | 2019 | Burrishoole | 0 | 0.67 | Whatman and Qiagen | Yellow | N/A | N/A | N/A |
| 63 | 2019 | Burrishoole | 6 | 0.67 | Whatman and Qiagen | Yellow | 64.5 | 465 | 5 |
| 64 | 2019 | Burrishoole | 13 | 0.85 | Whatman | Silver | 42,7 | 185 | 18.8 |
| 65 | 2019 | Burrishoole | 0 | 0.85 | Whatman | Silver | 38,5 | 105 | 22.7 |
| 66 | 2019 | Burrishoole | 16 | 0.85 | Whatman | Silver | 46,8 | 190 | 22 |
| 67 | 2019 | Burrishoole | 48 | 0.85 | Whatman | Silver | 50.7 | 215 | 20.2 |
| 68 | 2019 | Burrishoole | 15 | 0.85 | Whatman | Silver | 48.4 | 190 | 22.9 |
| 69 | 2019 | Burrishoole | 4 | 0.85 | Whatman | Silver | 51.6 | 210 | 22.9 |
| 70 | 2019 | Burrishoole | 2 | 0.85 | Whatman | Silver | 58.1 | 330 | 22.4 |
| 71 | 2019 | Burrishoole | 9 | 0.85 | Whatman | Silver | 31.5 | 50 | 22.5 |
| 72 | 2019 | Burrishoole | 5 | 0.85 | Whatman | Silver | 47.3 | 185 | 21.9 |
| 73 | 2019 | Burrishoole | 4 | 0.85 | Whatman | Silver | 51 | 230 | 23.3 |
| 74 | 2019 | Burrishoole | 6 | 0.85 | Whatman | Silver | 45.6 | 200 | 21.8 |
| 75 | 2019 | Burrishoole | 43 | 0.85 | Whatman | Silver | 51.2 | 215 | 19.2 |
| 76 | 2019 | Burrishoole | 0 | 0.85 | Whatman | Silver | 46.4 | 205 | 23 |
| 77 | 2019 | Burrishoole | 0 | 0.85 | Whatman | Silver | 43.6 | 165 | 27.6 |
| 78 | 2019 | Burrishoole | 13 | 0.85 | Whatman | Silver | 47.2 | 195 | 21.1 |
| 79 | 2019 | Burrishoole | 14 | 0.85 | Whatman | Silver | 47.4 | 190 | 24 |
| 80 | 2019 | Burrishoole | 15 | 0.85 | Whatman | Silver | 47.3 | 175 | 26.7 |
| 81 | 2019 | Burrishoole | 3 | 0.85 | Whatman | Silver | 40.9 | 130 | 24.3 |
| 82 | 2019 | Burrishoole | 3 | 0.85 | Whatman | Silver | 38.2 | 110 | 27.3 |
| 83 | 2019 | Burrishoole | 6 | 0.85 | Whatman | Silver | 49.7 | 250 | 24.3 |
| 84 | 2019 | Burrishoole | 14 | 0.85 | Whatman | Silver | 46.6 | 175 | 23.2 |
| 85 | 2019 | Burrishoole | 9 | 0.85 | Whatman | Silver | 42.4 | 145 | 25.6 |
| 86 | 2019 | Burrishoole | 0 | 0.85 | Whatman | Silver | 35.3 | 85 | 29.7 |
| 87 | 2019 | Burrishoole | 6 | 0.85 | Whatman | Silver | 44.2 | 175 | 20.7 |
| 88 | 2019 | Burrishoole | 10 | 0.85 | Whatman | Silver | 37.4 | 80 | 29.6 |
| 89 | 2019 | Burrishoole | 3 | 0.85 | Whatman | Silver | 38.8 | 100 | 26.5 |
| 90 | 2019 | Burrishoole | 46 | 0.85 | Whatman | Silver | 48.2 | 215 | 20.1 |
| 91 | 2018 | Lough Neagh | 3 | 0.68 | Qiagen | Yellow | 57.1 | 340 | N/A |
| 92 | 2018 | Lough Neagh | 0 | 0.68 | Qiagen | Yellow | 57.9 | 450 | N/A |
| 93 | 2018 | Lough Neagh | 0 | 0.68 | Qiagen | Yellow | 61.3 | 370 | N/A |
| 94 | 2018 | Lough Neagh | 1 | 0.68 | Qiagen | Yellow | 65.8 | 550 | N/A |
| 95 | 2018 | Lough Neagh | 4 | 0.68 | Qiagen | Yellow | 48.7 | 230 | N/A |
| 96 | 2018 | Lough Neagh | 4 | 0.68 | Qiagen | Yellow | 52.7 | 280 | N/A |
| 97 | 2018 | Lough Neagh | 3 | 0.68 | Qiagen | Yellow | 47.9 | 250 | N/A |
| 98 | 2018 | Lough Neagh | 4 | 0.68 | Qiagen | Yellow | 54 | 280 | N/A |
| 99 | 2018 | Lough Neagh | 5 | 0.68 | Qiagen | Yellow | 58.2 | 410 | N/A |
| 100 | 2018 | Lough Neagh | 4 | 0.68 | Qiagen | Yellow | 46.4 | 210 | N/A |
| 101 | 2018 | Lough Neagh | 0 | 0.68 | Qiagen | Yellow | 48.2 | 230 | N/A |
| 102 | 2018 | Lough Neagh | 0 | 0.68 | Qiagen | Yellow | 56.2 | 380 | N/A |
| 103 | 2018 | Lough Neagh | 1 | 0.68 | Qiagen | Yellow | 55.7 | 350 | N/A |
| 104 | 2018 | Lough Neagh | 1 | 0.68 | Qiagen | Yellow | 61.2 | 340 | N/A |
| 105 | 2018 | Lough Neagh | 0 | 0.68 | Qiagen | Yellow | 51.2 | 280 | N/A |
| 106 | 2018 | Lough Neagh | 0 | 0.68 | Qiagen | Yellow | 61.8 | 400 | N/A |
| 107 | 2018 | Lough Neagh | 0 | 0.68 | Qiagen | Yellow | 49.1 | 270 | N/A |
| 108 | 2018 | Lough Neagh | 1 | 0.68 | Qiagen | Yellow | 47.6 | 210 | N/A |
| 109 | 2018 | Lough Neagh | 4 | 0.68 | Qiagen | Yellow | 50.7 | 250 | N/A |
| 110 | 2018 | Lough Neagh | 0 | 0.68 | Qiagen | Yellow | 48.6 | 205 | N/A |
| 111 | 2018 | Lough Neagh | 2 | 0.68 | Qiagen | Yellow | 50.1 | 280 | N/A |
| 112 | 2018 | Lough Neagh | 0 | 0.68 | Qiagen | Yellow | 63.3 | 500 | N/A |
| 113 | 2018 | Lough Neagh | 7 | 0.68 | Qiagen | Yellow | 52.2 | 250 | N/A |
| 114 | 2018 | Lough Neagh | 1 | 0.68 | Qiagen | Yellow | 55.6 | 310 | N/A |
| 115 | 2018 | Lough Neagh | 4 | 0.68 | Qiagen | Yellow | 52.5 | 270 | N/A |
| 116 | 2018 | Lough Neagh | 3 | 0.68 | Qiagen | Yellow | 46.7 | 190 | N/A |
| 117 | 2018 | Lough Neagh | 2 | 0.68 | Qiagen | Yellow | 48.7 | 210 | N/A |
| 118 | 2018 | Lough Neagh | 14 | 0.68 | Qiagen | Yellow | 46 | 180 | N/A |
| 119 | 2018 | Lough Neagh | 11 | 0.68 | Qiagen | Yellow | 40.9 | 150 | N/A |
| 120 | 2018 | Lough Neagh | 0 | 0.68 | Qiagen | Yellow | 48.3 | 150 | N/A |
| 121 | 2018 | Lough Neagh | 6 | 0.68 | Qiagen | Yellow | 58.6 | 350 | N/A |
| 122 | 2018 | Lough Neagh | 12 | 0.68 | Qiagen | Yellow | 46.8 | 180 | N/A |
| 123 | 2018 | Lough Neagh | 9 | 0.68 | Qiagen | Yellow | 60.8 | 410 | N/A |
| 124 | 2018 | Lough Neagh | 0 | 0.68 | Qiagen | Yellow | 50 | 285 | N/A |
| 125 | 2018 | Lough Neagh | 1 | 0.68 | Qiagen | Yellow | 48.2 | 180 | N/A |
| 126 | 2018 | Lough Neagh | 1 | 0.68 | Qiagen | Yellow | 41.9 | 140 | N/A |
| 127 | 2018 | Lough Neagh | 0 | 0.68 | Qiagen | Yellow | 44.8 | 160 | N/A |
| 128 | 2018 | Lough Neagh | 4 | 0.68 | Qiagen | Yellow | 44.9 | 165 | N/A |
| 129 | 2018 | Lough Neagh | 0 | 0.68 | Qiagen | Yellow | 45.9 | 130 | N/A |
| 130 | 2018 | Lough Neagh | 3 | 0.68 | Qiagen | Yellow | 46.2 | 200 | N/A |
| 131 | 2018 | Lough Neagh | 6 | 0.68 | Qiagen | Yellow | 61 | N/A | N/A |
