## Supplementary 2 for "*In-situ* non-lethal rapid test to accurately detect the presence of the nematode parasite, *Anguillicoloides crassus*, in European eel, *Anguilla anguilla*"

**DNA Extraction from Whatman qualitative filter paper No. 1**

DNA Extraction

1. Prepare a 1cm^2^ cut of Whatman qualitative filter paper No. 1 in sterile condition

2. Deposit a drop of collected faecal material on the paper

3. Let air dry for approx. 5 min at room temperature

4. The piece can now be stored in -80◦C for long preservation or directly used to continue the extraction

5. Transfer each cut into a separate 1.5 ml Eppendorf and label with sample name

6. Add with 100 μl of FTA Purification Reagent

7. Incubate 5 min toom temperature

8. Remove the reagent and rinse with 200 μl of TE buffer

9. Remove carefully all TE buffer

10. Add 14 μl of Solution 1

11. Incubate at room temperature for 5 mins

12. Add 26 μl of Solution 2 to each Eppendorf

13. Incubate room temperature for 10 mins

14. Shake or votex for 10 sec

15. Centrifuge for 20 sec at 2000 rpm

16. Transfer 40 μl in a new sterile Eppendorf consisting in your final eluted DNA

Elution solutions

SOLUTION 1: 0.1N NaOH, 0.3mM EDTA, pH 13.0

-	Start with 50ml of stock 0.5M EDTA pH 8.0. Add solid NaOH slowly until you reach pH 13. Make	up to 250ml. This is now 0.1M EDTA pH13.

-	Make up 0.5M NaOH (20g of NaOH per 1 litre)

-	Add 200ml of 0.5M NaOH, 3ml 0.1M EDTA pH13, 797 ml H20.

SOLUTION 2: 0.1M Tris-HCl, pH7.0

-	Dissolve 12.1g of anhydrous Tris in less than a litre. pH using HCl to 7.0 and make up to a litre.
